# Haplotype resolved DNA methylome of African cassava genome

**DOI:** 10.1101/2022.09.12.507674

**Authors:** Zhenhui Zhong, Suhua Feng, Ben N. Mansfeld, Yunqing Ke, Weihong Qi, Yi-Wen Lim, Wilhelm Gruissem, Rebecca S. Bart, Steven E. Jacobsen

## Abstract

Cytosine DNA methylation is involved in biological processes such as transposable element (TE) silencing, imprinting, and X chromosome inactivation. Plant methylation is mediated by MET1 (mammalian DNMT1), DRM2 (mammalian DNMT3), and two plant-specific DNA methyltransferases, CMT2 and CMT3 (Law and Jacobsen, 2010). *De novo* DNA methylation in plants is established by DRM2 via the plant specific RNA-directed DNA methylation (RdDM) pathway that depends on two DNA-dependent RNA polymerases, Pol IV and Pol V (Gallego-Bartolome *et al*., 2019; Law and Jacobsen, 2010; Stroud *et al*., 2013). The DNA methylome of cassava has been previously documented based on its haploid collapsed genome (Wang *et al*., 2015). Since the cassava genome is highly heterozygous, DNA methylome analysis of the haplotype-collapsed genome misses many features of the methylome. With the development of long read sequencing and chromosomal conformation capture techniques, haplotype resolved genomes are available for highly heterozygous genomes (Mansfeld *et al*., 2021; Qi *et al*., 2022; Zhou *et al*., 2020), which provides high-quality reference genomes facilitating studies of haplotype resolved DNA methylomes.

---

To dissect the haplotype resolved DNA methylome of cassava, we conducted whole genome methylome studies in two haplotype genome resolved accessions of cassava (TME7 and TME204) using whole-genome bisulfite sequencing (WGBS) and enzymatic methyl-seq (EM-seq), respectively (Feng *et al*., 2020; Mansfeld *et al*., 2021; Qi *et al*., 2022). Sequencing reads were mapped to different haplotypes individually allowing zero mismatches and one best hit, which allowed the separation of reads belonging to different haplotypes. Overall, we found that the two haplotypes have similar whole genome methylation levels (Figure 1A, Supplemental Figure 1). We further plotted methylation levels over transcribed and flanking 1000 bp regions of protein-coding genes and transposable elements, and observed similar methylation levels between different haplotypes (Figure 1B,C, Supplemental Figure 2AB).

**Figure 1.**
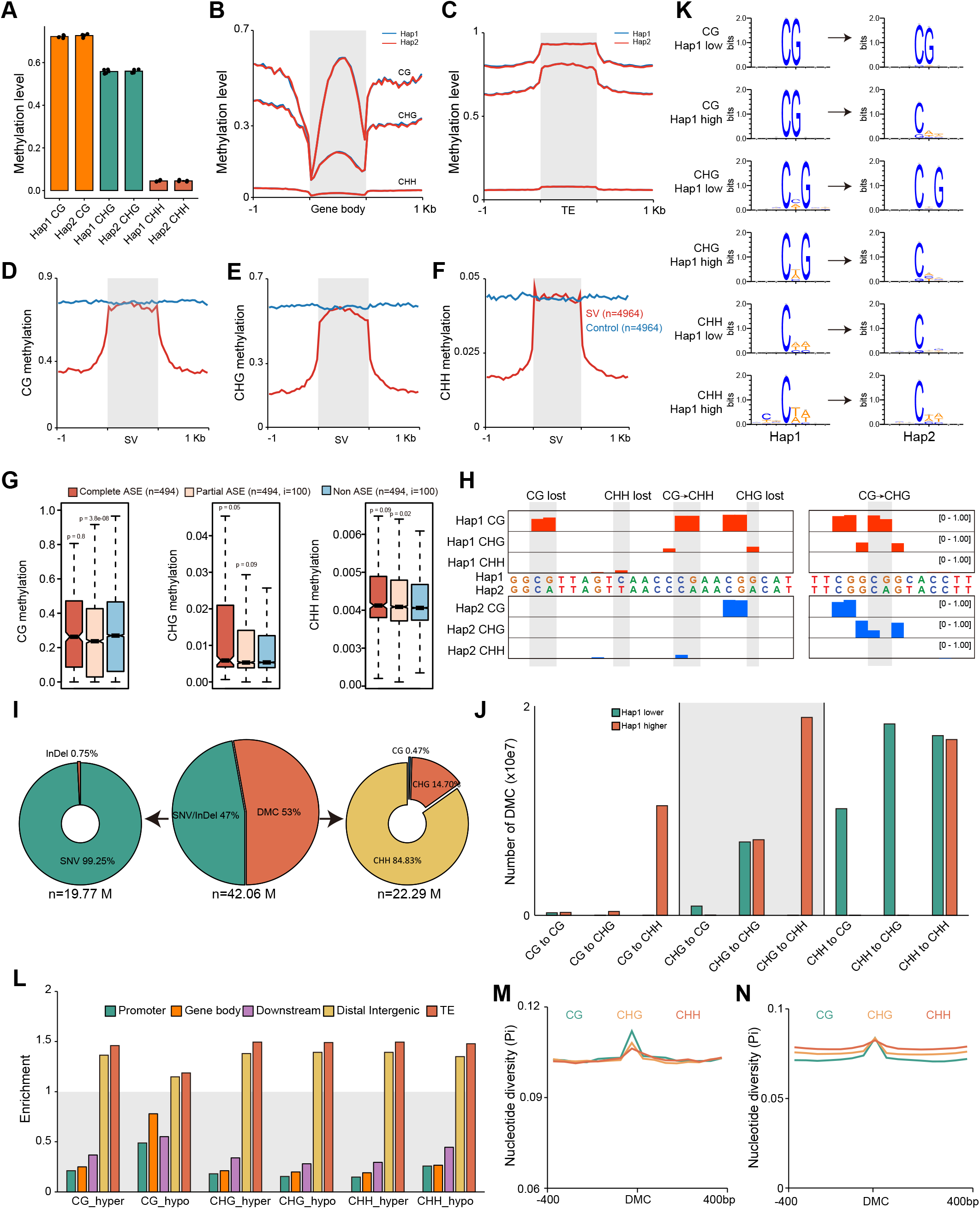
Haplotype resolved DNA methylome of African cassava. (A) Whole genome methylation levels of hap1 and hap2 haplotypes in TME7 (n=3) and TME204 (n=1). (B) Metaplot of CG, CHG, and CHH methylation levels over protein-coding genes and flanking 1Kb sequences of hap1 and hap2 haplotypes in TME204. (C) Metaplot of methylation levels over transposon elements and flanking 1Kb sequences in TME204. (D-F) Metaplot of methylation levels over structural variation regions and flanking 1Kb sequences in TME7. Equal length regions were randomly selected in the genome (blue line). (G) Methylation levels of complete, partial, and non-ASE genes in TME7. (H) Representative screenshots of DMCs. Methylation site losses and mC context changes are indicated on top of the track (red: hap1; blue: hap2). (I) Ratio of DMCs caused by SNP/InDel, where the cytosines are lost, and DMCs caused by different methylation levels. (J) Numbers of different types of cytosine context variations between the two haplotypes. (K) Consensus sequences of DMCs. (L) Genomic distribution enrichment of DMCs. (M-N) Nucleotide diversity of 400bp flanking regions of DMC in TME7 (M) and TME204 (N). Panels I, J, L, and K show results for TME7. Results for TME204 are shown at Supplemental Figure 2.

Previous studies have revealed large numbers of haplotype-specific structural variants (SVs) in cassava (Mansfeld *et al*., 2021; Qi *et al*., 2022). To understand how DNA methylation is associated with these SVs, we analyzed methylation levels of these SVs. Focusing on places where haplotype-specific SVs occur, we found that flanking regions of SVs have approximately two times lower methylation levels than random controls and SVs, suggesting that SVs preferentially take place at lowly methylated regions, with the SVs turning these regions to heavily methylated regions (Figure 1D-F). Allele-specific expression (ASE) refers to preferential expression of the allele transcribed from one haplotype (Gaur *et al*., 2013). ASEs can be cataloged into complete ASEs and partial ASEs. Since the number of partial ASEs and non-ASE controls are much higher than that of complete ASEs, we selected an equal number of complete ASEs and partial ASEs or non-ASE controls with 100 times iterations. Interestingly, complete ASE genes showed higher gene body CHG methylation than partial (p = 0.05) and non-ASE genes and partial ASE genes showed lower CG methylation than others (p = 3.8e-08, Figure 1G).

Next, we analyzed differentially methylated cytosines between haplotypes (haplotypic DMCs). We first aligned the two haplotypes and identified syntenic cytosines (Supplemental Figure 2C). Methylation levels of syntenic cytosines were compared between two haplotypes. We observed at least three scenarios (Figure 1H) that resulted in differential methylations between haplotypes: 1) SNP/InDel in one haplotype leads to the loss of the cytosine (47.17%) among which more than 99.25% are SNP variants (Figure 1I; see TME204 data in Supplemental Figure 2D); 2) Cytosine context alterations (e.g. CG in hap1 and CHH in hap2) that leads to the methylation level changes detected in our analysis (28.98%); and 3) Cytosines stay in the same contexts between haplotypes but exhibit different methylation levels (23.85%). For DMCs caused by scenarios 2 and 3, the majority of them were CHH variations (Figure 1I). When CG sites are mutated into CHG or CHH sites, they are more likely to be hypomethylated; whereas CHH sites are more likely to be hypermethylated when mutated into CHG and CG sites (Figure 1JK, Supplemental Figure 2EF). Furthermore, we found that DMCs are frequently detected at TEs and distal intergenic regions, while depleted at gene body and flanking regions, suggesting that TEs are hot spots for frequent DNA methylation changes (Figure 1L, Supplemental Figure 2G). Finally, we investigated the genetic diversity of DMC sites and 400 bp flanking sequences and found that nucleotide diversity of DMC sites are significantly higher than that of flanking sequences (Figure 1MN), demonstrating that DMC sites are under more frequent natural selection. Together, our analyses compared haplotype resolved DNA methylomes of cassava, and place genomic heterozygosity within the haplotypic epigenetic regulatory landscape.

## Acknowledgements

This research was supported by grants from the Bill & Melinda Gates Foundation (OPP1194889). S.E.J. is an Investigator of the Howard Hughes Medical Institute.

## Authors’ contributions

Z.Z., S.F., B.M., W.G., R.B. and S.J. conceived and designed the research. Z.Z., W.Q., and Y.K. performed data analysis. Y.L. and S.F. prepared sequencing libraries. Z.Z., S.F., and S.J. wrote the manuscript.

## Data availability statement

The DNA methylome data are available in Gene Expression Omnibus under accession number GSE192748 (https://www.ncbi.nlm.nih.gov/geo/query/acc.cgi?acc=GSE192748).

