## Supplementary figures for "Haplotype resolved DNA methylome of African cassava genome"

### Supporting Information

### Method

#### DNA methylation library preparation

Methylome of TME7 samples were sequenced by whole genome bisulfite sequencing (WGBS). Briefly, genomic DNA was end-repaired and ligated with TruSeq DNA single-end adapters (Illumina) using a Kapa DNA HyperPrep kit (Roche). Adapter-ligated DNA was converted with an EpiTect Bisulfite Kit (Qiagen). Converted DNA was PCR-amplified by MyTaq polymerase (Bioline) for 12 cycles. Methylome of TME204 sample was sequenced by Enzymatic Methyl-seq (EM-seq). EM-seq libraries were prepared from sheared DNA using an Enzymatic Methyl-seq kit following manufacturer instructions (New England BioLabs) with 6 cycles of PCR (Feng et al., 2020). The libraries were run on a D1000 ScreenTape (Agilent) to determine the quality and size, and then purified by AMPure XP beads (Beckman Coulter). Library concentrations were measured with a Qubit dsDNA Broad-Range Assay kit (ThermoFisher). Libraries were sequenced on a HiSeq 2500 or NovaSeq 6000 sequencer (Illumina).

#### Methylome mapping

WGBS and EM-seq reads were mapped to haplotype 1 and haplotype 2 genomes of TME7 (Mansfeld et al., 2021) or TME204 (Qi et al., 2022) by Bsmap (v2.90) allowing 0 mismatches and 1 best hit (-v 0 -w 1) (Xi & Li, 2009). Duplicated reads were removed with SAMtools (v1.3.1) (Li et al., 2009). Reads with three or more consecutive methylated CHH sites were considered as unconverted reads and removed in the following analysis. Conversion rate was estimated by calculating methylation level of the chloroplast genome. DNA methylation level at each cytosine was calculated by number of methylated C vs. total C and T account. To call Differentially Methylated Cytosines (DMCs), two haplotype genomes were aligned with each other by LastZ with script published previously (Zhou et al., 2020). Only cytosines with at least 3000 bp syntenic flanking regions were kept in the analysis. Two-tailed Fisher’s exact test was used to calculate p-value between syntenic cytosines. We used differences in CG,

CHG, and CHH methylation > 0.4, 0.2, and 0.1, respectively, and p-value < 0.05 as cutoff to define DMCs. Methylation track files were visualized with Integrative Genomics Viewer (IGV, v3.0) (Robinson et al., 2011). Structural Variations (SVs) and Allele Specific Expressions (ASEs) information are from previous studies (Mansfeld et al., 2021; Qi et al., 2022).

**Consensus sequence, DMC enrichment, and nucleotide diversity analysis**

Sequence consensus analysis was performed with WebLogo (v3.6.0) (Crooks et al., 2004). Flanking 4 bp sequences of DMC were obtained with bedtools (v2.26.0) getfasta function and used as input sequences of WebLogo (Quinlan & Hall, 2010). Genomic distribution enrichment analysis was conducted with ChIPseeker and bedtools (Yu et al., 2015). DMC sites and random control sites of equal number were used as input of ChIPseeker. For transposable element (TE) enrichment analysis, DMC sites and random control sites of equal number were intersected with TE regions to calculate sites that distributed within or outside TE regions. For nucleotide diversity analysis, DMC sites and flanking 400 bp regions were used as input of Pim (v0.3) with 50 bp window length (-w) and 50 bp steps between windows (-S) (Haubold et al., 2011).

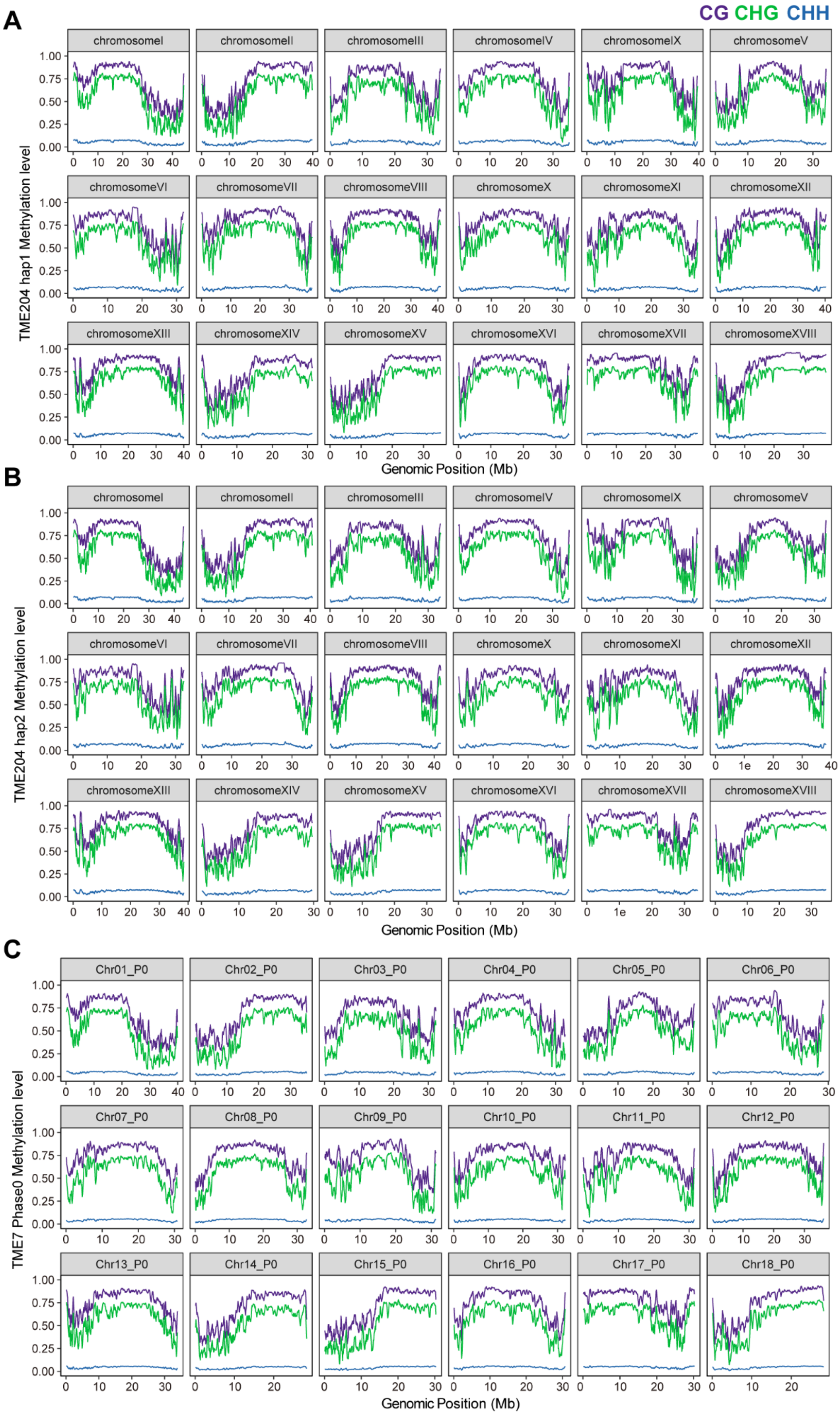


**Supplemental Figure 1. Whole genome methylation over chromosomes of TME204 and TME7.**

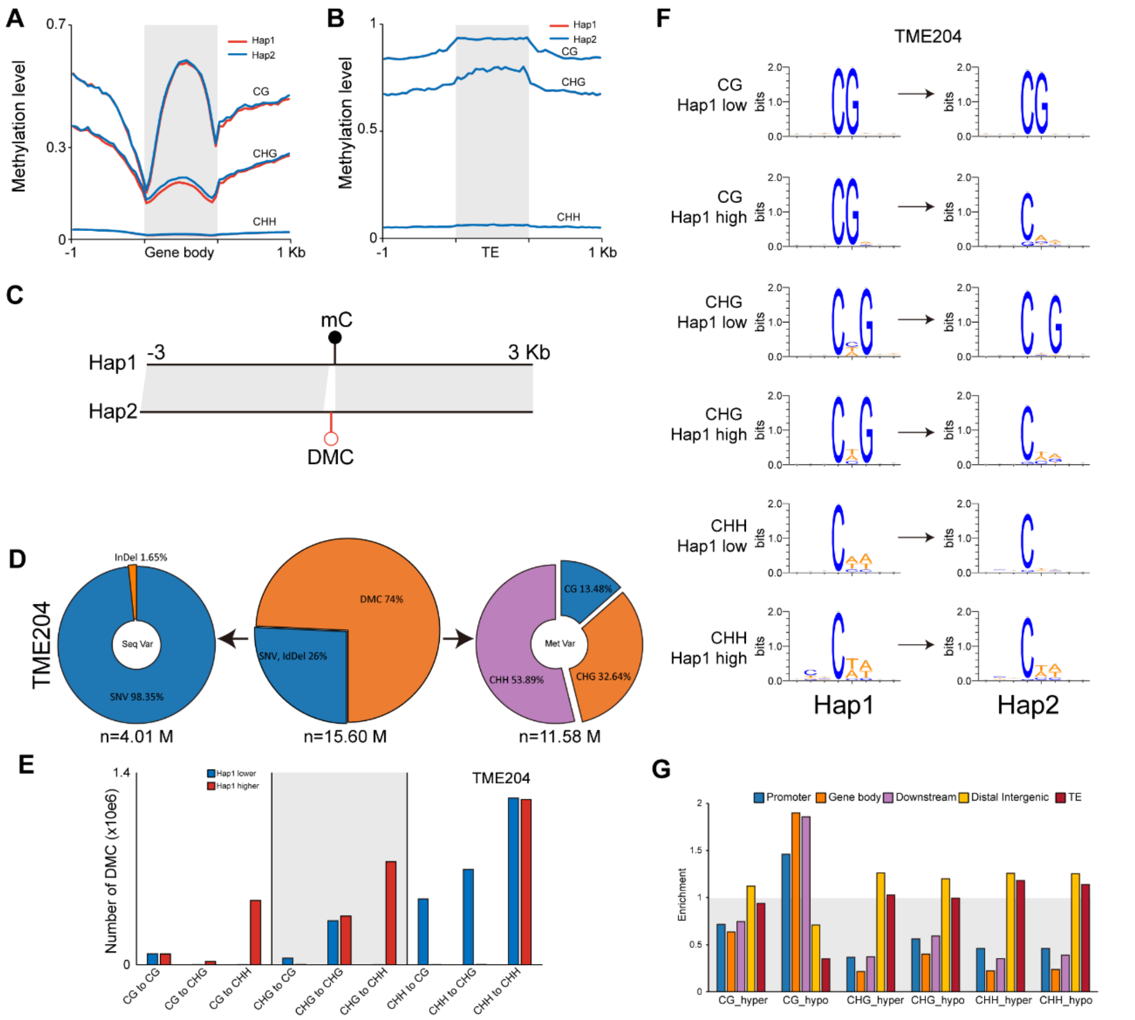


**Supplemental Figure 2 Haplotype resolved DNA methylome of African cassava.** (A) Metaplot of CG, CHG, and CHH methylation levels over protein-coding genes and flanking 1Kb sequences of hap1 and hap2 haplotypes in TME7. (B) Metaplot of CG, CHG, and CHH methylation levels over transposon elements and flanking 1Kb sequences in TME7. (B) Metaplot of CG, CHG, and CHH methylation levels over transposon elements and flanking 1Kb sequences in TME7. (C) Schematic diagram showing identification of differentially methylated cytosine (DMC). Only cytosines with > 3Kb flanking sequences aligned between the two haplotypes are analyzed. (D) Ratio of DMCs caused by SNP/InDel, where the cytosines are lost, and DMCs caused by different methylation levels (differences in CG > 0.4, in CHG > 0.2, and in CHH > 0.1) in TME204. (E) Numbers of different types of cytosine context variations between the two haplotypes of TME204. (F) Consensus sequences of DMCs of TME204. (G) Genomic distribution enrichment of DMCs of TME204.
